# Genetic and Biochemical Characterization Reveals PduU as a Modular Tool for Tuning Pdu Microcompartment Permeability

**DOI:** 10.64898/2026.07.29.741414

**Authors:** Komal S. Timane, Chiranjit Chowdhury

## Abstract

Bacterial microcompartments (MCPs) are versatile proteinaceous organelles that compartmentalize metabolic pathways, offering promising scaffolds for synthetic biology and metabolic engineering. However, designing customized nanobioreactors requires distinguishing structurally indispensable shell proteins from those that can be modified or deleted to tune shell permeability without disrupting core organelle assembly. In this study, we performed a systematic biophysical and metabolic characterization of the hexameric shell protein PduU to evaluate its potential as a modular platform for synthetic organelle engineering. We tested whether deleting *pduU* or selectively truncating its N-terminal β*-barrel* domain preserves shell assembly, metabolite flux, and intermediate confinement. Our results demonstrate that PduU modifications alter shell permeability while fully maintaining organelle structural integrity, monodispersity, and electrostatic colloidal stability. Crucially, this modulation in permeability redirects internal metabolic flux toward the energy-generating propionate pathway, resulting in elevated cell biomass and significantly increased yields of propionate, an economically vital industrial platform chemical. By establishing that PduU is a non-essential structural component whose modification tunes small-molecule flux, this work highlights PduU as a flexible locus for shell engineering, providing a scalable strategy for biomanufacturing of high-value bio-based products in tailor-made MCP nanobioreactors.

## Introduction

MCPs are protein-bounded metabolic organelles that spatially organize enzymatic pathways to enhance reaction efficiency while preventing the escape of volatile or toxic intermediates. One of the most extensively studied MCP systems is the 1,2-propanediol utilization (Pdu) microcompartment of *Salmonella enterica*, which enables coenzyme B_12_-dependent degradation of 1,2-propanediol while sequestering the volatile intermediate propionaldehyde within a selectively permeable protein shell. Structural and biochemical studies over the past two decades have established that this shell is composed of multiple paralogous proteins that assemble into hexamers, pseudohexamers, and pentamers to form a pseudo-icosahedral enclosure [1], [2].

Early structural work demonstrated that canonical shell proteins possess central pores consistent with metabolite transport. For example, PduA forms a wide, polar pore compatible with substrate influx, while PduT contains a metal-centered pore housing a [4Fe−4S] cluster associated with redox balancing or cofactor exchange [2], [3]. These studies established a foundational principle in MCP biology: pore architecture in shell proteins correlates directly with specific transport functions. However, not all shell components conform to this paradigm. Several low-abundance components of the Pdu shell exhibit architectures that do not obviously support passive metabolite diffusion. This observation raises broader questions recognized across MCP biology: how do specialized shell proteins contribute to functions beyond simple permeability, and how is shell composition linked to metabolic performance?

A striking structural outlier in the Pdu shell is PduU. Unlike canonical single-domain hexamers (PduA/J) or tandem-domain trimers (PduB/T), the crystal structure of PduU revealed a circularly permuted BMC fold in which the canonical pore position is occluded by a six-stranded N-terminal β-barrel, whose interior is tightly packed with bulky side chains incompatible with passive metabolite diffusion [4]. On the opposite face of the hexamer, PduU presents a large hydrophobic cavity lined with aromatic residues -a feature absent from classical pore-forming shell proteins [4]. Based on these unique structural hallmarks, it was originally suggested that PduU possesses a “special functional role” distinct from transport, although no functional tests of this proposal were performed [4].

Genetic analysis of the Pdu operon provided an important physiological context for interpreting this structure. Systematic gene deletions in *S. enterica* demonstrated that major shell proteins (PduA, PduB/B′, PduJ, and PduN) are essential for MCP structural integrity and aldehyde sequestration, whereas deletion of *pduU* (*del pduU*) disrupts neither overall MCP morphology nor intermediate confinement [5]. Instead, the *del pduU* mutant exhibits a significant reduction in growth rate during growth on 1,2-propanediol under coenzyme B_12_-replete conditions while behaving similarly to the wild type under B_12_ limitation [5]. This phenotype differs fundamentally from structural shell mutants that display leaky shells and altered permeability, indicating that PduU is dispensable for building the shell superstructure but is required for optimal metabolic output during high pathway flux [5].

A systems-level perspective on MCP organization further strengthens this view. Interactome modeling combined with bacterial two-hybrid validation identified a direct interaction between PduU and the cytosolic GTPase PduV, suggesting that the prominent β-barrel region of PduU may serve as an outward-facing interaction interface between the MCP and the cytoplasm [6]. However, the specific residues involved, the structural basis of this interaction, and its precise physiological relevance remain uncharacterized. Recent conceptual advances in MCP biology provide a modern framework for interpreting these observations. Reviews of MCP shell architecture now emphasize that shell proteins are not static structural tiles, but dynamic, polymorphic modules capable of specialized roles in permeability, assembly, spatial alignment, and metabolic regulation [7], [8]. Within this framework, low-stoichiometry shell components with unusual folds are increasingly viewed as regulatory or organizational elements rather than passive structural units [7], [8].

Taken together, structural, genetic, physiological, and interactome data converge on a consistent but unresolved picture: PduU is structurally incompatible with passive diffusion, dispensable for shell assembly, evolutionarily conserved, and physiologically important under conditions of high pathway flux, yet its molecular function remains unknown [2], [4], [6]. Importantly, none of the prior studies directly tested whether the distinctive β-barrel and hydrophobic cavity of PduU contribute to metabolite exchange, protein interaction, or internal MCP organization [4].

In this study, we evaluate whether PduU and its unique β-barrel capping domain can be engineered or exploited as a customizable platform for synthetic nanoreactor design. To systematically dissect its functional role and shell tolerance, we constructed knockouts of the full *pduU* gene (*del pduU*) and a targeted β-barrel deletion variant (*del N-ter pduU*), subsequently evaluating their growth kinetics under both B_12_-saturating and B_12_-limiting conditions using 0.4% 1,2-propanediol as the primary carbon source. To confirm locus-specific phenotypes, plasmid-based ectopic expression was performed for genetic complementation. At the organelle level, we purified intact microcompartments from these variants and comprehensively analyzed their structural integrity and biophysical properties through SDS-PAGE, transmission electron microscopy (TEM), dynamic light scattering (DLS), and zeta potential measurements. Furthermore, we assessed how these shell modifications influence luminal enzymatic activity and quantified overall metabolite transport kinetics across the shell via HPLC. By combining genetic, structural, and metabolic characterization, this work reveals how PduU influences internal catalytic performance, providing critical insight into its suitability as a robust, non-essential locus for engineering functionalized synthetic nanobioreactors.

## Material and Methods

### Chemicals and reagents

1,2- Propane diol, vitamin B_12_ (CN- B_12_), glycerol, 3-Methyl-2-BenzoThiazolinone Hydrazone hydrochloride (MBTH), NAD+, NADH, 4-(2-aminoethyl) benzenesulfonyl fluoride hydrochloride (AEBSF) and alcohol dehydrogenase (from *Saccharomyces cerevisiae*), Lysozyme, Coenzyme A were from Sigma-Aldrich (St. Louis, MO). Antibiotics, Isopropyl-β-D- 1-thiogalactopyranoside (IPTG), dithiothreitol (DTT) and arabinose were from Himedia (Mumbai, India). Q5 DNA polymerase, restriction enzymes, and HiFi DNA Assembly Master Mix were from New England Biolabs (Beverly, MA). NovaTaq PCR Master Mix was from Merck (Darmstadt,Germany). 4-chloro- dl- phenylalanine, n-Octyl-β-D-thioglucopyranoside (OTG) and other chemicals were obtained from Qualigens Thermofisher Scientific (Powai, India).

### Bacterial strains, media and growth conditions

The bacterial strains used in this study are listed in Table S1. All strains are derivatives of *Salmonella enterica* serovar Typhimurium strain LT2. The rich medium used was Luria broth (Lennox) (LB) medium (Himedia, Mumbai, India) (G. Bertani, 1951). Tryptone yeast extract (TYE) medium was used for the selection of transformants where bacto-tryptone, yeast extract, and agar were purchased from Himedia Laboratories (Himedia, Mumbai, India). The minimal medium used was no carbon-E (NCE) medium supplemented with 1 mM MgSO4, 0.3 mM each of valine, isoleucine, leucine and threonine, and 50 μM Ferric citrate [5], [9]. Growth studies were performed at limiting (20 nM) and saturating (100 nM) CN-B_12_ concentrations, which serve as exogenous B_12_ source. The growth was monitored as previously described using BioTek Epoch 2 Microplate Spectrophotometer (Agilent-BioTek, India) [5], [10].

### Three-dimensional model building and visualization

The three- dimensional structure of PduU β- barrel mutant was modelled using the Swiss Model server [11] with PDBID 3CGI as a template and all the visualization and graphical presentations were carried out in Pymol [12].

### Construction of chromosomal mutations and complementation strains

All primers used in this study are listed in Table S2. Chromosomal deletion of the gene encoding PduU shell protein (*del pduU*) and its β- barrel region (*del N-ter pduU*) were constructed by recombineering as follows [3], [5], [13]. Chromosomal modifications were introduced using lambda RED-mediated homologous recombination via plasmid pKD46. A dual-selectable *mpheS-kan* cassette (Kanamycin resistance / counter-selectable *mpheS*) was amplified with 40 bp target-homology flanks. The linear PCR product was introduced by electroporation into the lambda-RED expressing host strain. Recombinants were selected on Kanamycin plates and verified for correct genomic integration via colony PCR and sequencing. Synthetic DNA containing specific mutations (*pduU* gene and β –barrel region of *pduU* gene deletions) was electroporated into the cassette-containing strain. Transformants were counter-selected on media containing DL-4-chlorophenylalanine (which inhibits strains expressing *mpheS*). Resistant colonies were screened for Kanamycin and ampicillin sensitivity to confirm both the loss of the *mpheS-kan* cassette and the curing of the temperature-sensitive pKD46 plasmid. Final mutant constructs were confirmed by PCR amplification and DNA sequencing. For complementation studies, the genes encoding PduU shell protein was cloned into the pLac22 plasmid within the *Bgl*II and *Hin*dIII sites [5] and confirmed by PCR followed by sequencing.

### MCP purification, diol dehydratase (DDH) and propionaldehyde dehydrogenase (PduP) activity assays

Pdu microcompartments were purified by differential centrifugation following detergent-assisted lysis [1], [14]. Briefly, induced *S. enterica* cells were lysed using a buffer containing lysozyme, protease inhibitors (AEBSF), and mild detergent (1.5% OTG). Cell debris was pelleted by low-speed centrifugation (12,000×g), and intact MCPs were subsequently isolated from the supernatant by high-speed centrifugation (20,000×g), washed, and resuspended in storage buffer for downstream analysis.

Luminal enzymatic activities were determined spectrophotometrically at 340 nm following published protocols [1], [15]. Diol dehydratase (DDH) activity was measured via a coupled assay in which the propionaldehyde product was converted to 1-propanol by yeast alcohol dehydrogenase, and the continuous rate of NADH oxidation was monitored. Propionaldehyde dehydrogenase (PduP) activity was measured by monitoring the rate of NAD+ reduction to NADH in a reaction mixture containing propionaldehyde, Coenzyme A HS-CoA, and NAD+.

### Dynamic Light Scattering and Electron microscopy

The hydrodynamic diameter (particle size distribution) and surface charge (zeta-potential) of purified Pdu microcompartments were measured using a Zetasizer Nano instrument (Malvern Panalytical,) at 25^°^C according to established electrophoretic and dynamic light scattering protocols [16], [17]. Purified MCP samples were diluted in filtered (0.22μm) storage buffer to avoid multiple scattering effects and particle aggregation. All measurements were performed in triplicate (n = 3), and results are reported as the mean ± standard deviation (SD).

For electron microscopy purified MCPs were negatively stained with uranylless and visualized using a transmission electron microscope (JEOL JEM F200) as described earlier [1], [18].

### Propionaldehyde Quantification Assay

Propionaldehyde accumulated in the growth medium was quantified using a 3-methyl-2- benzothiazolinone-hydrazone hydrochloride (MBTH) assay adapted to a 48-well microplate format. Culture samples (40 μL) were periodically collected from baffled flasks in a fixed time interval (every 4 h). The samples were diluted as necessary and added to microplate wells containing 100 mM potassium citrate buffer (pH 3.6), 0.1% (w/v) MBTH solution, and double-deionized water to bring the initial reaction volume to 715 μL.

The microplates were incubated at 37°C for 15 min to allow complete derivatization of propionaldehyde by MBTH. Following incubation, 285 μL of double-deionized water was added to each well (yielding a final volume of 1 mL), and the absorbance was measured at 305 nm using a microplate reader. Propionaldehyde concentrations were calculated using a standard curve generated with known propionaldehyde standards (0.01- 0.2 mM linear range) [19].

### High-Performance Liquid Chromatography (HPLC) quantification of metabolites

Extracellular concentrations of the substrate (1,2-propanediol) and metabolic end-products (1- propanol and propionate) in wild-type and mutant culture supernatants were quantified using High-Performance Liquid Chromatography (HPLC), adapted from previously described methods [1], [20]. Aliquots 1 mL of culture media were collected at every 4 h time interval; bacterial cells were cleared off by centrifugation (5000 rpm for 5 min) followed by filtered through 0.22 μm nylon membrane filters. Separation was performed using an organic acid analysis column (Aminex HPX-87H, 300 x 7.8, Bio-Rad). Isocratic elution was carried out using 5 mM H_2_SO_4_ as the mobile phase at a constant flow rate of 0.4 mL/min. Metabolites were detected using a Refractive Index (RI) detector. Compounds were identified by comparing retention times to those of authentic pure standards (1,2-propanediol, 1-propanol, and propionate), and concentrations were calculated using linear calibration curves constructed for each standard.

### Statistical analysis

Where appropriate, statistical analysis was completed to determine the significance of observed differences using two-tailed unpaired *t*-tests with a significance level of 0.05. Differences between two experimental groups were analyzed using a two-tailed unpaired Student’s *t*-test. Statistical significance was defined as p < 0.05. All statistical analyses were conducted using [GraphPad Prism v8.0].

## Results

### Targeted deletion of PduU and its N-terminal β-barrel domain extends lag phase without decreasing doubling time or terminal cell density

Under B_12_-limiting conditions (20nM), both mutant strains displayed a prominent lag phase delay compared to WT which is approximately 25-30h post-inoculation. Despite the extended lag phase, once exponential growth was initiated, the doubling times of the mutants were comparable to or slightly faster than WT (Td = 12.07 ± 1.18h): *del pduU*: Td = 10.2 ± 0.63h: *del N-ter pduU* Td = 11.43 ± 0.83h. Both mutant strains achieved equivalent or slightly higher final optical densities (OD_600_ approximately 0.53 to 0.56 relative to WT OD_600_ 0.49 at 48 h (Fig 1A and C).

**Fig 1:**
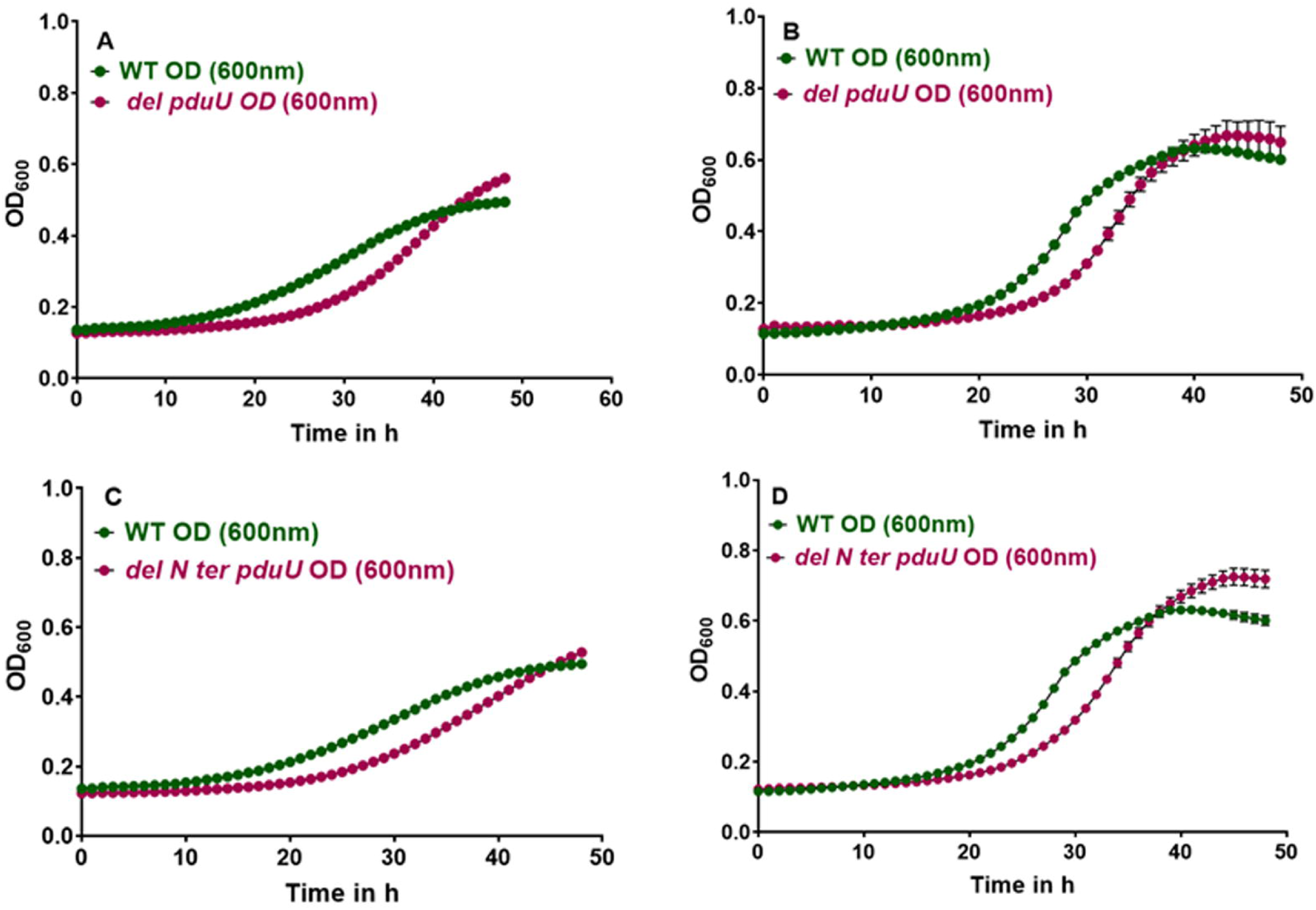
Representative Graphs for growth (at 37 degrees) of wild type *Salmonella enterica* Serovar Typhimurium LT2, and *del pduU* (1A and B) and *del N-ter pduU* (1C and D) on minimal medium supplemented with 0.4% 1,2 propanediol as carbon source with limiting B_12_(20 nM) (A and C) and saturating B_12_(100 nM) (B and D) concentrations. The error bars represent one standard deviation and are based on three biological replicates.

When supplied with saturating coenzyme B_12_ (100 nM), overall metabolic flux increased for all strains, resulting in a roughly twofold reduction in doubling times across all the strains. Similar to the limiting condition, both *del pduU* and *del N-ter pduU* mutants exhibited a delayed entry into log phase (lagging behind WT by roughly 5-8 h). Exponential growth rates were nearly identical between WT (Td = 5.94 ± 0.29 h) and the mutant strains: *del pduU* Td = 5.5 ± 0.17h: *del N-ter pduU* Td = 6.44 ± 0.22 h. Notably, both mutant strains eventually surpassed WT in terminal optical density under 100 nM B_12_ (Fig 1B and D).

### Ectopic expression of PduU restores normal growth kinetics in complementation assays

To confirm that the extended lag phase observed in the deletion strains was locus-specific and not the result of polar effects or secondary mutations, plasmid-based complementation assays were performed. Expressing *pduU* in trans from the isopropyl-beta-D-1-thiogalactopyranoside (IPTG)-inducible vector pLAC22 (pLAC22-*pduU*) in both the mutant backgrounds completely rescued the growth phenotype upon induction with 500 μM IPTG (Fig S1A and B).

### PduU and its **β**-barrel domain are dispensable for microcompartment assembly and core protein composition

To determine whether the deletion of *pduU* or its N-terminal β-barrel domain affects organelle assembly or protein encapsulation, microcompartments were purified from WT and mutant strains. SDS-PAGE analysis (Fig 2A) and subsequent densitometric analysis (Fig 2B) showed no significant alterations in major proteins bands compared to wild-type MCPs. Major shell proteins (such as PduA, PduB/B’, PduJ, and PduN) alongside key luminal enzymatic machinery (PduCDE, PduP) were retained (Fig 2A). This confirms that microcompartments maintain full assembly capacity and efficient internal enzyme loading even in the complete absence of PduU or its N-terminal β-barrel domain.

**Fig 2:**
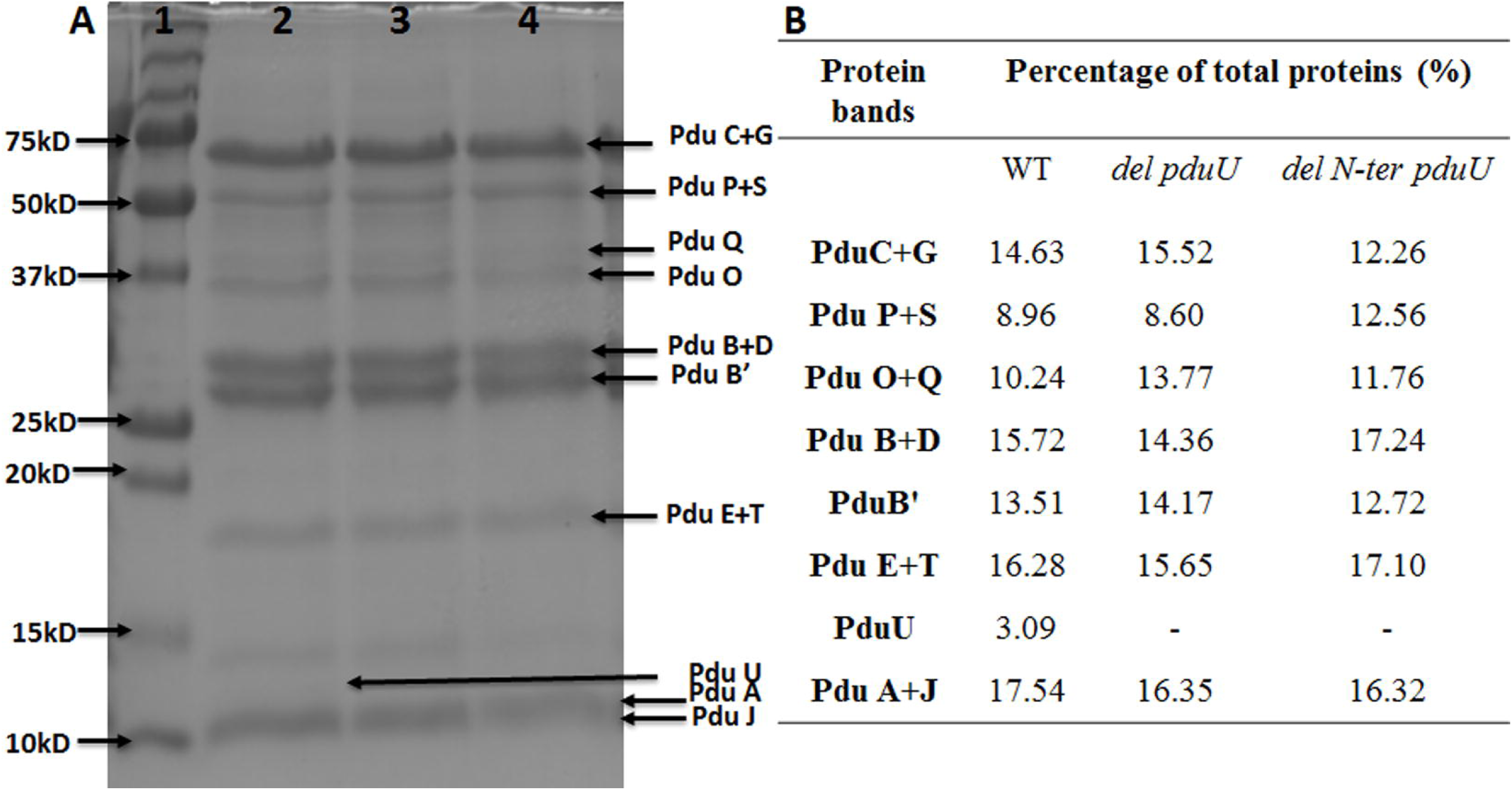
A:SDS PAGE of WT and mutant MCPs. Lane1- Molecular weight ladder (kDa), Lane 2- WT MCP, lane 3- *del pduU* MCP and lane 4- *del N-ter pduU* MCP. Equal amounts of protein (20μg) were loaded per lane on 15% SDS PAGE. B: Densitometric analysis of WT and mutant MCPs. Relative protein abundance was determined by densitometric analysis using ImageJ v2.16.0, with individual band intensities expressed as a percentage of total MCP protein (the sum of all resolved bands). Overlapping bands were quantified jointly, such as PduP+S representing the combined contribution of both proteins. Densitometry demonstrated a linear response between 5 and 25 µg of loaded protein. Representative values were obtained by resolving ∼20 µg of purified MCPs on a 4–12% SDS-PAGE gel at 150 V for 1 h, followed by Coomassie Brilliant Blue staining

The isolated MCP yield per gram of wet cell biomass was calculated and WT baseline yield was determined to be 1.41-1.87 mg MCP/g cells (100%). While overall protein composition was unchanged, both *del pduU* and *del N-ter pduU* showed a modest to moderate increase in recovery, yielding 108%-147% (2 mg MCP/g cells for *del pduU* and 2.38 mg MCP/g cells for *del N-ter pduU*) compared to WT.

### Biophysical characterization reveals preserved polyhedral architecture with distinct surface modulations

To evaluate the physical dimensions, structural morphology, and surface charge properties of purified organelles, transmission electron microscopy (TEM), dynamic light scattering (DLS), and electrokinetic surface potential (zeta-potential) measurements were performed. Transmission electron micrographs confirmed that purified MCPs from all three strains (WT, *del pduU*, and *del N-ter pduU*) maintained intact, classic polyhedral shell geometries without visual evidence of structural collapse or destabilization (Fig 3A). Complementary DLS analysis (Fig 3B) demonstrated that WT and *del pduU* MCPs exhibit virtually identical hydrodynamic diameters and narrow size distributions (Zaverage = 147.37 ± 3.48 nm, PDI = 0.12 ± 0.04 for WT; Zaverage = 148.20 ±7.21 nm, PDI = 0.14 ± 0.05 for *del pduU*). In contrast, the *del N-ter pduU* displayed a modest increase in hydrodynamic diameter (204.30 ± 10.96 nm while preserving a low polydispersity index (PDI = 0.100 ± 0.026) (Table 1).

**Fig 3:**
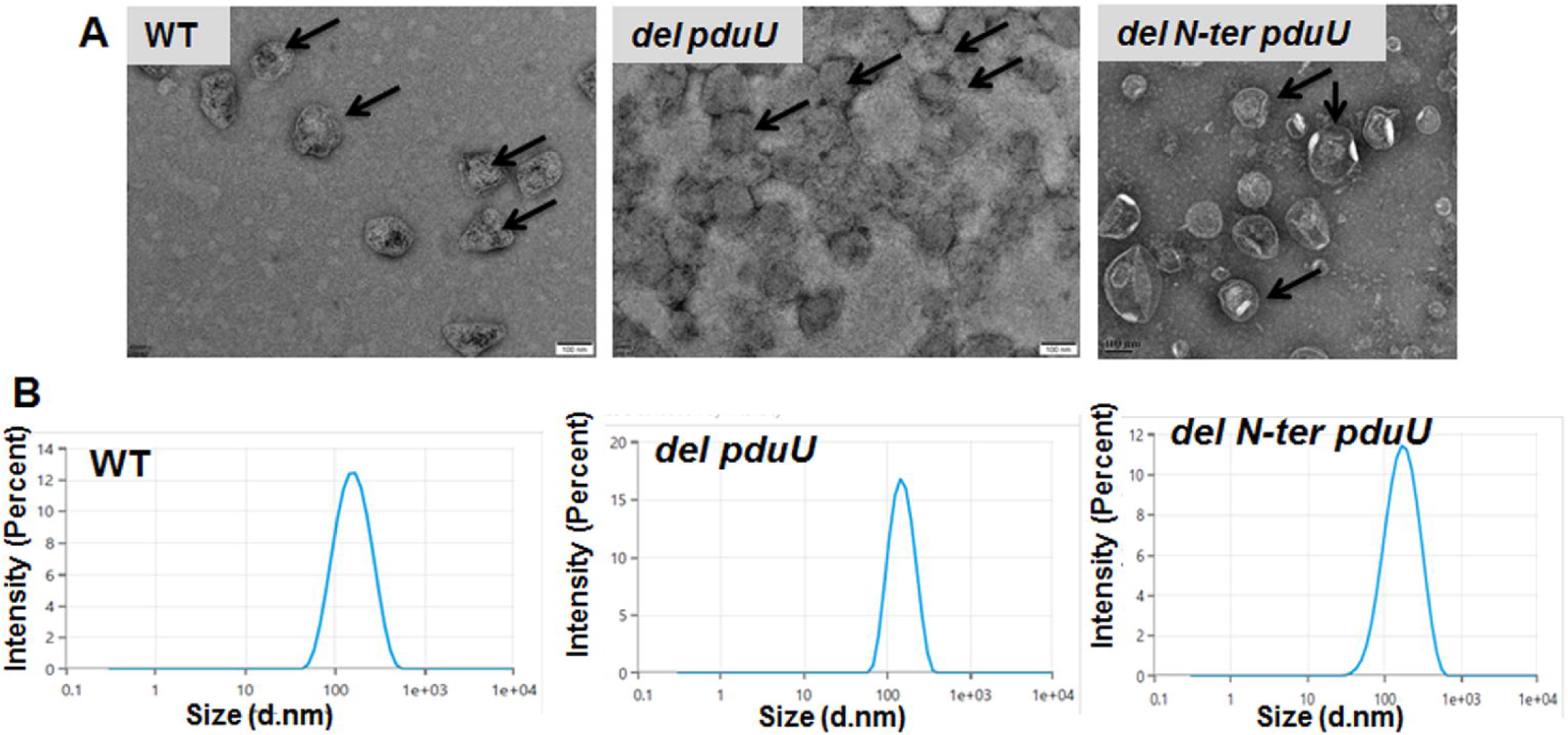
A: Transmission electron micrographs of purified wild-type and mutant PduU microcompartments. B: Representative intensity-weighted size distribution profile of wild-type and mutants measured by Dynamic Light Scattering (DLS) at 25 °C.

**Table 1:**
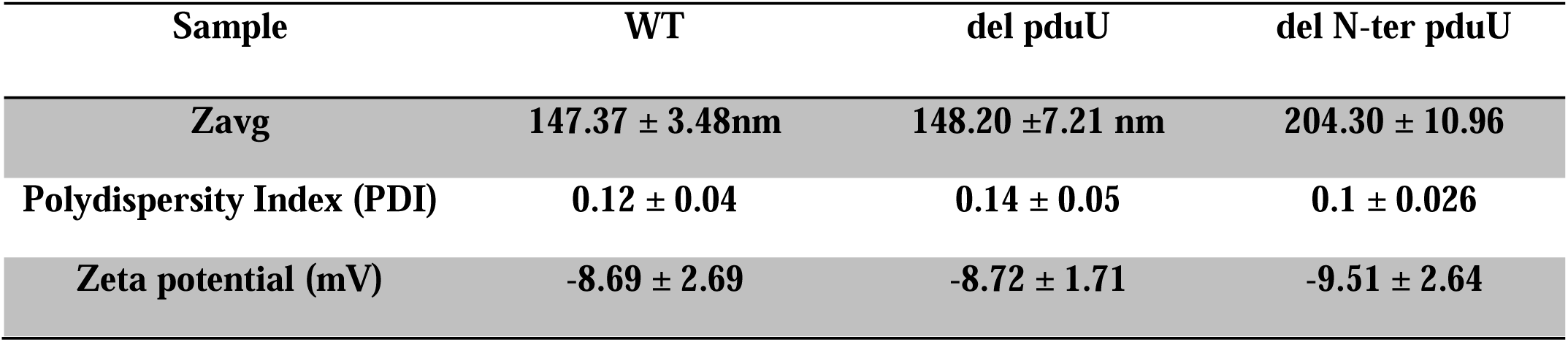
The Dynamic light scattering and zeta potential of WT and mutants.

Electrokinetic surface charge analysis revealed that all purified MCPs carry a net negative surface potential in storage buffer, with the mutant strains exhibiting a progressive shift toward higher negative magnitude (zeta = −8.69 ± 2.69 mV for WT; zeta = −8.72 ± 1.71 mV for *del pduU*; zeta = −9.51 ± 2.64 mV for *del N-ter pduU*) (Table 1).

### Luminal Diol Dehydratase and Propionaldehyde Dehydrogenase activities remain substantially intact in shell-modified microcompartments

To determine whether deletion of PduU or its N-terminal β-barrel domain impairs the catalytic functionality of encapsulated enzymes, the relative *in-vitro* enzymatic activities of diol dehydratase (DDH,) and propionaldehyde dehydrogenase (PduP) were measured from intact, purified MCPs and normalized to WT. These enzymatic results directly align with our SDS-PAGE and biophysical analyses. First, the retention of robust relative DDH and PduP activities (93%–96%) confirms that neither *pduU* deletion nor truncation of its β*-barrel* domain interferes with the correct folding, assembly, or encapsulation of core luminal enzymes during microcompartment biogenesis (Fig 4).

**Fig 4:**
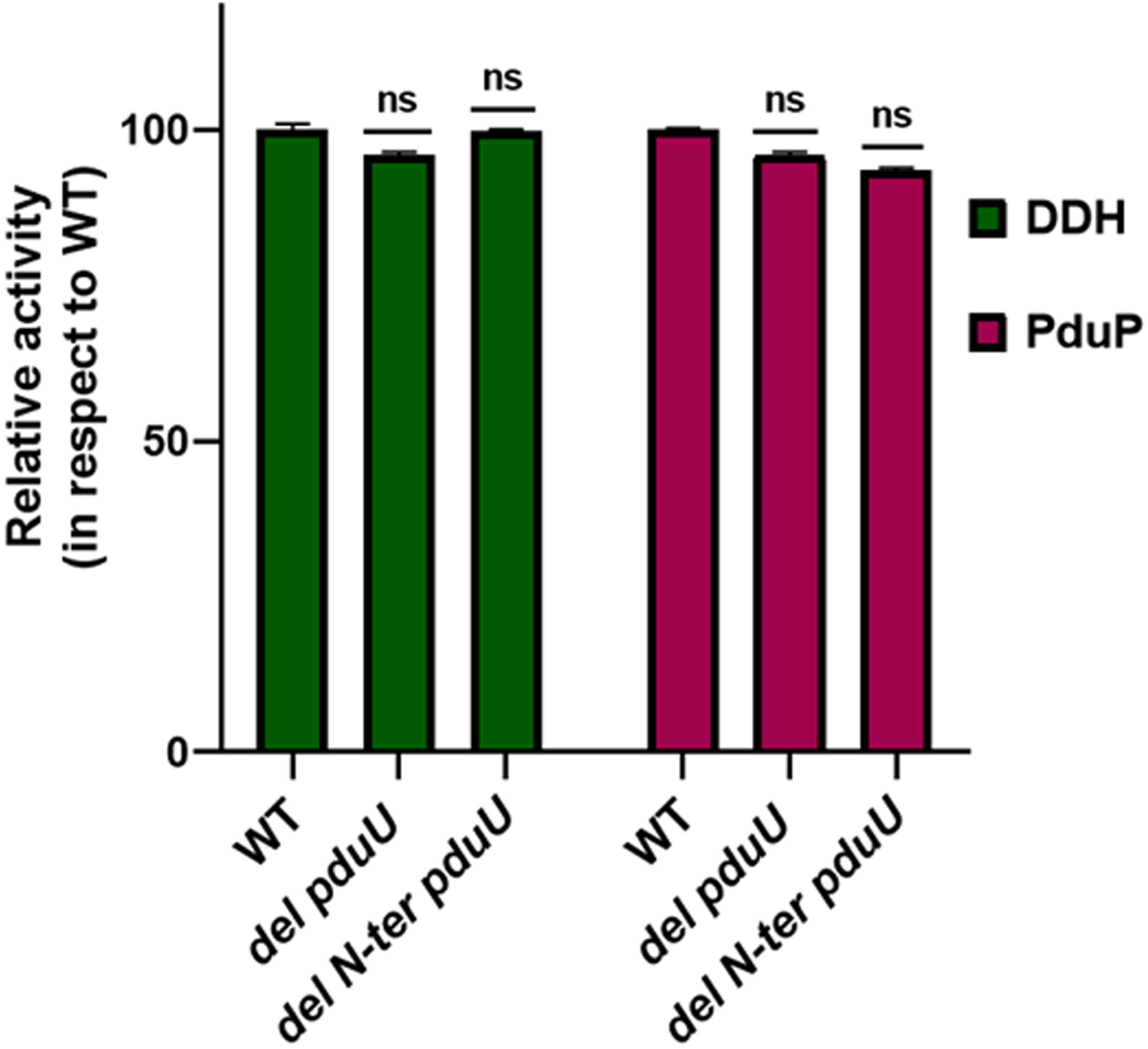
Relative DDH and PduP enzyme activities of wild-type and mutant MCPs. Activities are presented as percentages relative to the WT control (set to 100%). Data represent mean ± SD from three independent biological replicates (n = 3).

### HPLC metabolite profiling reveals delayed substrate depletion and shifted end-product kinetics

To evaluate how shell structural modifications alter metabolic flux across the organelle boundary in living cells, HPLC time-course profiling was conducted to quantify extracellular substrate (1, 2-propanediol), and end-products (propionate and 1-propanol) in culture supernatants over 48 h.

In WT, 1,2-PD consumption initiated rapidly (0-12 h) and was completely exhausted by 24 h. In both mutants, 1,2-PD depletion exhibited a distinct kinetic lag, remaining largely unchanged during the initial 0-16 h. Once active consumption began (16-28 h), substrate depletion proceeded rapidly; reaching total consumption by 28 h (Fig 5A and B). The generation of 1- propanol exhibited a similar 4-6 h kinetic delay in both mutant strains, reaching a maximum concentration (approximately 16-18 mM) between 24-28h before clearance by 44h (Fig 6A and B).

**Fig 5:**
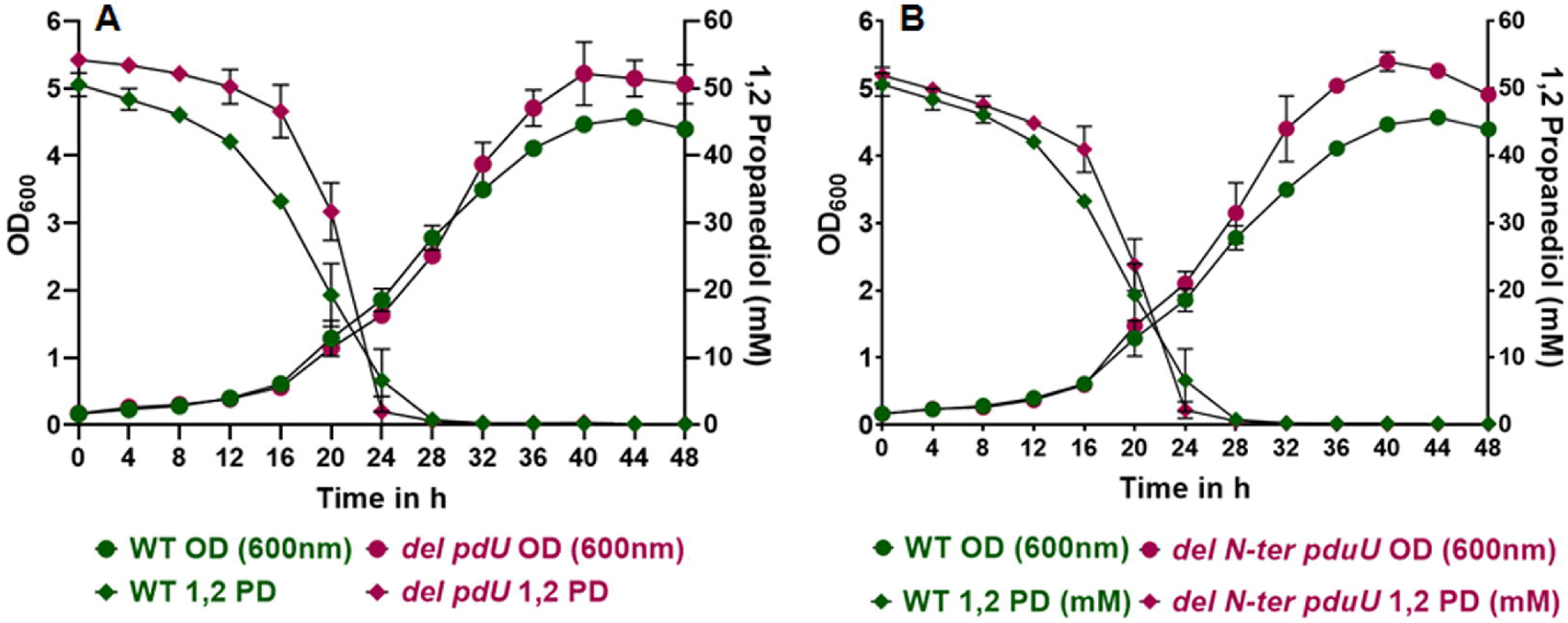
Effects of mutants on substrate uptake in aerobic growth measured by the HPLC. Cells were grown in baffled flasks and samples were taken at fixed time interval. Data is presented as mean ± SEM from three independent biological experiments (n = 3).

**Fig 6:**
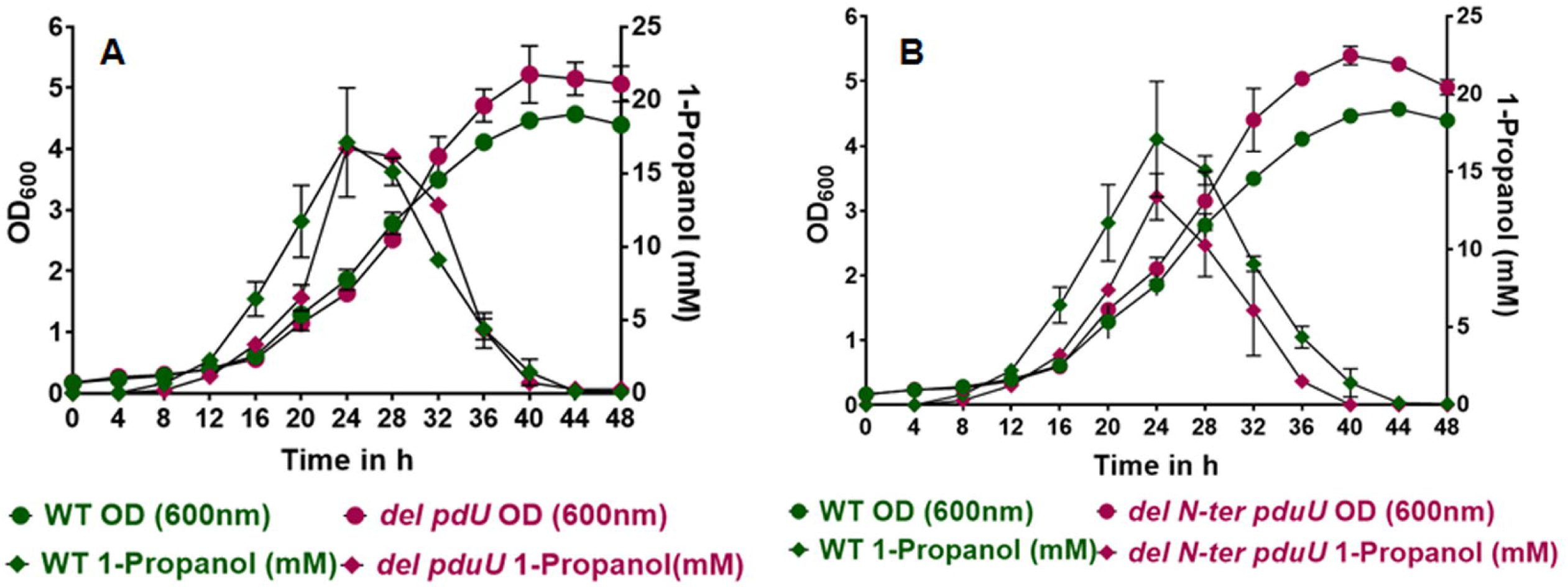
Effects of mutants on product formation (1-propanol) in aerobic growth measured by the HPLC. Cells were grown in baffled flasks and samples were taken at fixed time interval. Data are presented as mean ± SEM from three independent biological experiments (n = 3).

A notable outcome of *pduU* disruption is a substantial increase in peak extracellular propionate concentration relative to WT. While WT cultures reached a maximal propionate accumulation of approximately 7.0 mM at 24 h, the *del pduU* and *del N-ter pduU* mutants accumulated peak levels of approximately 10.5 mM (+40%) and approximately 8.5 mM (+21%), respectively (Fig 7A and B).

**Fig 7:**
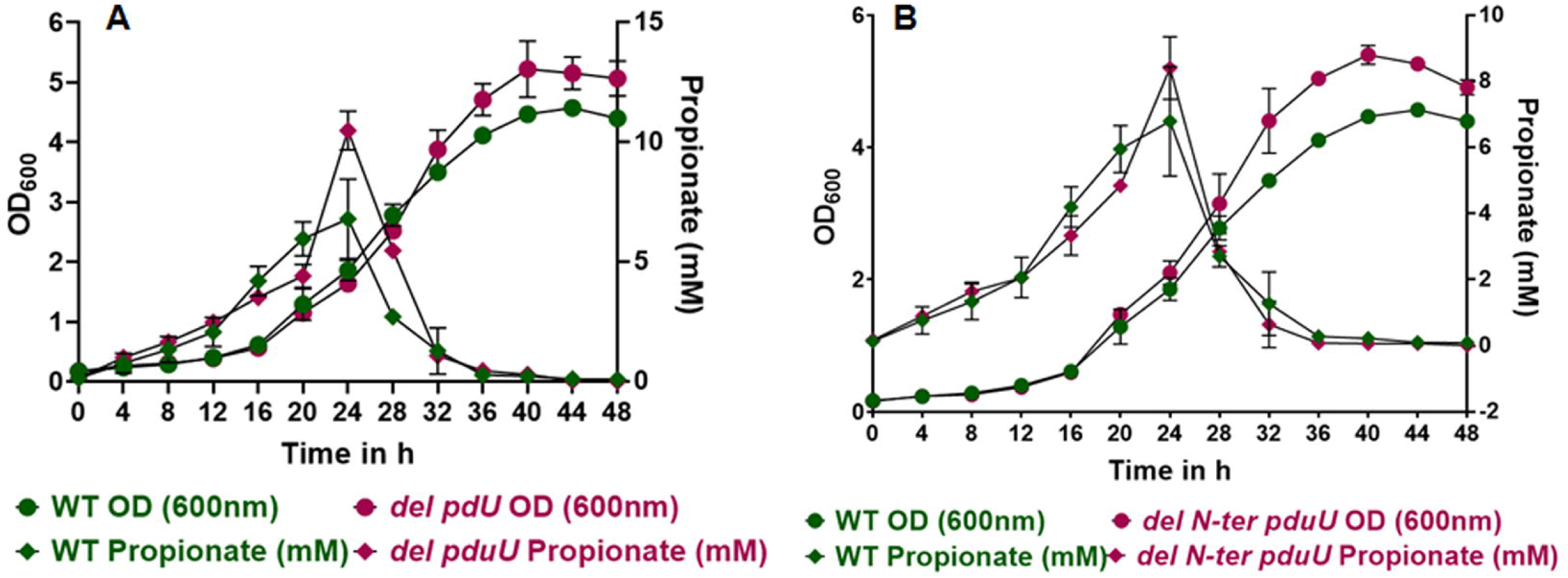
Effects of mutants on product formation (propionate) in aerobic growth measured by the HPLC. Cells were grown in baffled flasks and samples were taken at fixed time interval. Data are presented as mean ± SEM from three independent biological experiments (n = 3)

### Propionaldehyde Quantification

In WT, propionaldehyde levels peaked modestly at 20 h (approximately 1.5 mM) and were rapidly cleared as biomass increased. In both mutants, extracellular propionaldehyde reached significantly higher peak levels (approximately 2.5 mM at 24 h for *del pduU*; 1.7 mM at 16 h for *del N-ter pduU*) before being completely metabolized by 36 h (Fig 8A and B).

**Fig 8:**
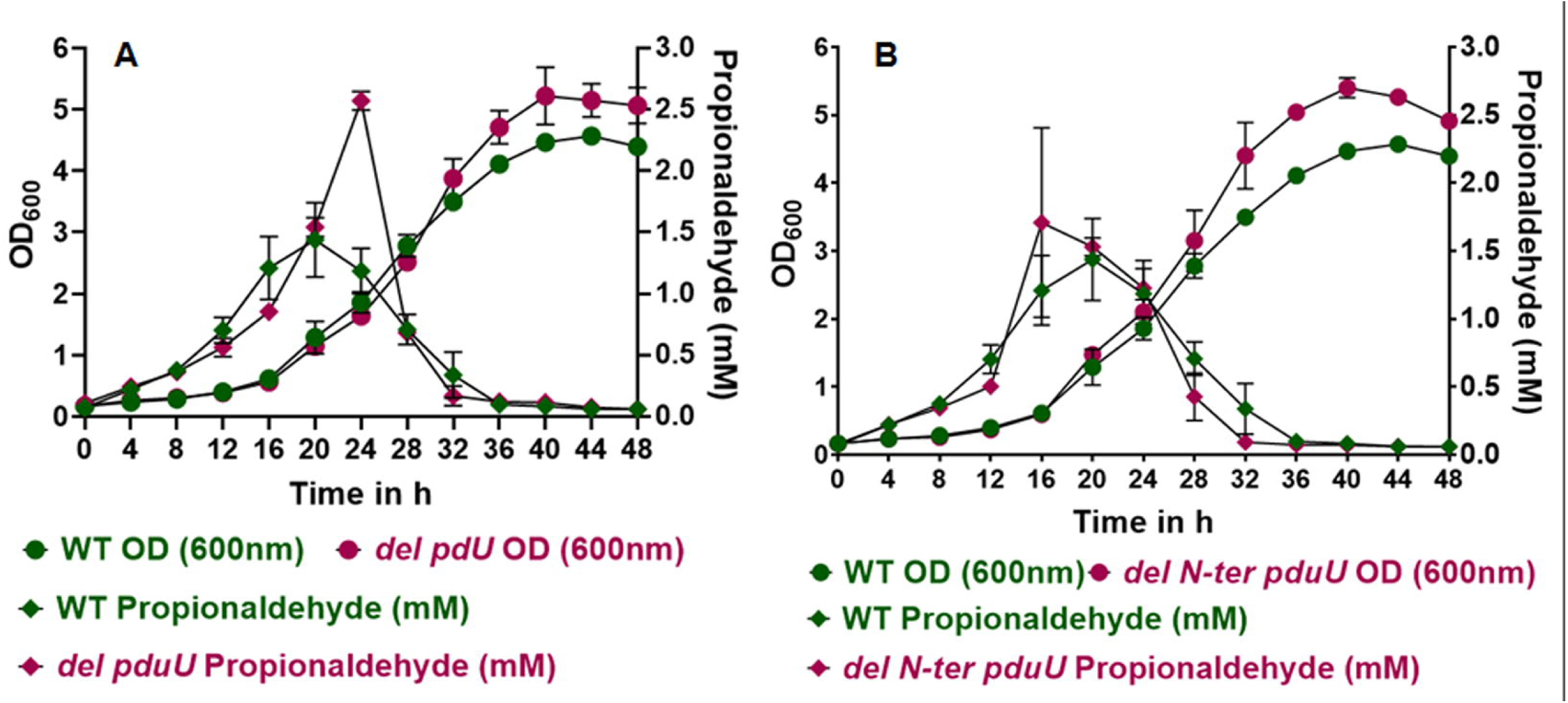
Effects of mutants on propionaldehyde formation in aerobic growth. The propionaldehyde was quantified using a 3-methyl-2-benzothiazolinone-hydrazone hydrochloride (MBTH) assay by derivatization of propionaldehyde with MBTH and measuring the absorbance of the complex at 305 nm in microplate reader. Data is presented as mean ± SEM from three independent biological experiments (n = 3)

## Discussion

MCPs are complex, self-assembling proteinaceous organelles that spatial-localize specialized metabolic pathways. Understanding the specific structural roles of individual shell components, particularly low-abundant proteins with unusual structural folds is essential for deciphering native organelle biology and advancing rational synthetic bio-engineering [8], [21]. In this study, we systematically characterized the hexameric shell protein PduU its prominent N-terminal β- barrel.

The growth phenotypes under limiting (20 nM) and saturating (100 nM) coenzyme B_12_ conditions are consistent with the earlier findings [5]. In that PduU is non-essential for 1,2- propanediol utilization. However, these earlier studies reported a general reduction in exponential growth rate, our growth profiling indicates that the disruption of *pduU*, specifically its N-terminal β-barrel domain primarily induces a kinetic delay during early pathway activation (extended lag phase), without compromising the exponential doubling rate or final biomass yield once log-phase growth is established.

Physical characterization confirmed that microcompartment architecture remains remarkably resilient to PduU modifications. Dynamic Light Scattering (DLS) revealed near-identical hydrodynamic diameters for WT and *del pduU* MCPs, alongside a modest increase for the *del N-ter pduU* variant. All preparations maintained narrow, highly uniform particle populations with exceptionally low polydispersity indices (PDI ≤ 0.100). In nanoparticle and colloidal chemistry, a PDI < 0.30 strictly denotes a highly monodisperse, uniform, and homogeneous population [16], [22]. These consistently low PDI values confirm that neither deleting *PduU* nor removing its N-terminal β-barrel domain compromises structural uniformity or causes non-specific organelle aggregation.

Furthermore, electrokinetic analysis showed that all MCPs retain a net negative surface potential. This net negative surface charge generates critical mutual electrostatic repulsion between neighboring organelles, preventing self-aggregation and confirming that deleting *PduU* or its N-terminal β*-barrel* domain reinforces particle colloidal stability in buffer systems used in the study.

Since DDH is a signature enzyme of Pdu MCP and operates entirely within the internal lumen, the persistence of near-native reaction rates in intact organelles demonstrates that deleting *pduU* or truncating its N-terminal β-barrel domain does not restrict the influx of the primary substrate (1,2-propanediol) across the shell boundary and confirm that the modified protein shell remains fully permeable to substrate diffusion. Instead, the primary consequence of PduU disruption is a subtle alteration in intermediate (propionaldehyde) retention. Time-course propionaldehyde quantification revealed that modifying PduU makes the shell slightly more permeable, resulting in a temporal increase in extracellular propionaldehyde leakage levels.

This temporary loss of intermediate leakage initially imposes metabolic stress and slows internal flux, presumably triggering the observed kinetic lag phase. To compensate for this initial bottleneck, the cells might strategically reallocate the metabolic flux toward the energy-generating propionate pathway mediated by PduP and downstream enzymes. This metabolic shift is evidenced by a substantial increase in propionate accumulation by +40% in the *del pduU* 21% in *del N-ter pduU* variant, compared to the WT. By prioritizing propionate synthesis over 1- propanol reduction during substrate processing, the cell yields additional ATP to overcome the early intermediate leakage stress. Ultimately, this energetic gain allows the mutants to resume robust growth and achieve higher final biomass yields upon entering stationary phase.

The previous interactome modeling and bacterial two-hybrid assays originally suggested that PduU connects the microcompartment to the cytoplasm via a direct interaction with the cytosolic GTPase PduV [6]. Based on this, it could be hypothesized that the phenotypic lag and intermediate leakage observed in *del pduU* stem indirectly from mislocalizing PduV away from the organelle shell. However, our *vivo* experimental evidence disproves this model: a *del pduV* deletion strain exhibits wild-type growth kinetics on 1,2-propanediol (Fig S2A), and ectopic expression of *pduV* (*del pduU* / pLAC22-*pduV*) upto 1 mM IPTG fails to rescue the *del pduU* phenotype (Fig S2B). Together, these results confirm that the metabolic disruption in *del pduU* is an intrinsic consequence of altering shell permeability, rather than a secondary loss of PduV recruitment.

Further, 3-D modeling analysis of N-terminal β-barrel domain truncated PduU variant revealed a large open central pore. The pore is capable of facilitating the unhindered flux of propionate across the microcompartment shell (Fig S3).

## Conclusion

Over the past decade, MCPs have emerged as versatile, highly modular platforms for synthetic biology and metabolic engineering, offering a natural solution for compartmentalizing complex or toxic metabolic pathways [23]. Because MCP shells are inherently amenable to genetic and structural modifications, researchers have successfully repurposed these proteinaceous architectures into custom bio-nanoreactors [24], [25], [26]. These established applications highlight the immense potential of engineered MCPs as scalable bio-manufacturing scaffolds [8], [27]. A major hurdle in synthetic nanobioreactor development is determining which shell components are structurally essential versus those amenable to modification without compromising organelle integrity. Here, we investigated whether deleting or truncating the low abundant shell protein PduU maintains shell assembly, metabolic flux, and intermediate confinement.

In nature, native microcompartment systems, such as the metabolosome identified in *Clostridium phytofermentans*, encapsulate specialized catabolic pathways that ferment plant cell wall-derived deoxyhexose sugars (such as fucose and rhamnose) through a 1,2-propanediol intermediate into major products including propionate, propanol, and ethanol [28].

Driving higher yields of propionate by tuning the 1,2 ropanediol utilization microcompartment permeability holds immense industrial relevance. Propionate (propionic acid) is a high-value platform chemical widely utilized across the food, feed, pharmaceutical, and chemical industries [29]. Due to its powerful antimicrobial and antifungal properties, it serves as a critical, safe preservative in bakery products and animal feed. Furthermore, propionate is an essential bio-based building block for synthesizing non-steroidal anti-inflammatory drugs, specialty solvents, and sustainable bioplastics like cellulose acetate propionate [30]. Because commercial propionate is still predominantly derived from petroleum-based feedstocks through energy-intensive thermochemical processes, engineering microbial platforms such as shell-modified microcompartments for elevated propionate flux provides a sustainable, bio-based alternative for industrial-scale manufacturing.

In conclusion, targeting non-essential shell features like the PduU β-barrel offers a powerful, rational design tool. By balancing isolation yields, stable structural integrity, and tunable pore transport, these engineered PduU variants represent high-capacity modular tool nanobioreactor engineering, industrial biomanufacturing, and targeted enzymatic biocatalysis.

## Acknowledgements

KST acknowledges the fellowship awarded by the Department of Biotechnology (Award No- DBT/JRF/BET-18/i/2018/AL/245-1064), Ministry of Science and Technology, Government of India. CC acknowledges the Science and Engineering Research Board (SERB), Government of India (CRG/2019/001558) and Indian Council of Medical Research (Diarr/Adhoc/5/2022-ECD-II) for financial support. Authors also acknowledge Council of Scientific and Industrial Research (CSIR) - National Chemical Laboratory, Pune, India for academic support.

## Declaration of Competing Interest

Authors declared that there is no known competing conflict of interest with respect of publication of this review article.

## Declaration of generative AI and AI-assisted technologies in the writing process

During manuscript preparation, OpenAI was utilized to assist with grammatical editing and language polish. The author(s) subsequently reviewed, verified, and revised all generated text, and assume full accountability for the scientific accuracy, claims, and overall content of the final article.

## Notes

### Competing Interest Statement

The authors have declared no competing interest.

### Summary of Updates

The title has been changed, supporting information has been updated, and typos have been corrected.

