## Supplementary for "Genetic and Biochemical Characterization Reveals PduU as a Modular Tool for Tuning Pdu Microcompartment Permeability"

**Tables**

**Table S1**. Bacterial strains used in the study

| **Species/Strain** | **Genotype** | **Source** |
| --- | --- | --- |
| ***E. coli*** | | |
| **YS9** | DH5alpha/pKD46 | Lab Collection |
| **MB1** | DH5alpha/pLac22 |  |
| **CE30** | *E.coli* top 10/pJD141 |  |
| ***Salmonella enterica* Typhimurium LT2** | | |
| **CE1** | LT2 *Salmonella enterica* serovar Typhimurium LT2 | Lab collection |
| **CE19** | Δ*pduV*::frt |  |
| **KST18** | ∆*pduU* | This Study |
| **KST41** | ∆*N-ter pduU* |  |
| **KST23** | ∆*pduU/*pLac22 |  |
| **KST29** | ∆*pduU*/pLac22-*pduU* |  |
| **KST66** | ∆*pduU*/pLac22-*pduV* |  |
| **KST101** | Δ*N-ter pduU*/pLac22 |  |
| **KST104** | ∆*N-ter pduU*/pLac22-*pduU* |  |

**Table S2**. Primers used in this study

| **Description** | **Primer** | **Sequence (5’-3’)** | |
| --- | --- | --- | --- |
| **Mutant Preparation** |  |  |  |
| ***pduU*** | *pduU- mphe-F* | | TGATCCCACGCCCGCATGAAGCCATGTGGCGACAGATGGTGGAGGGGTAA GAATTCGCGGCCGCATCTAG |
|  | *pduU-mphe-R* | | AAACATCAAACGCTTCATGACTTTACGTCCGGGTGATCGAGCAAGTGGTG CTGCAGCGGCCGCTACTAGT |
| ***N-ter pduU*** | *Barrel pduU-mphe-F* | | GACAGATGGTGGAGGGGTAATGGAAAGACAACCGACAACGGAATTCGCGGCCGCATCTAG |
|  | *Barrel pduU-mphe-R* | | GATTAGCAATCAGGTGCGCGAGAGTGACCTGTTTCCCCGGCTGCAGCGGCCGCTACTAGT |
| **Clone preparation** |  |  |  |
| ***pduU* clone in pLac22** | *pduU*-Gib-PLac-F | | TAACAATTTCACACAGGAAAGATCTATGGAAAGACAACCGACAAC |
|  | *pduU*-Gib-PLac-R | | GTGATAAACTACCGCATTAAAGCTTTTACGTCCGGGTGATCGAG |
| **Mutant and clone confirmation** |  |  |  |
| **pLac22** | PLac22-F | | CCCCAGGCTTTACACTTT |
|  | PLac22-R | | GTTAGATTTCATACACGGTGCCTG |
| ***pduU*** | *pduU-F* | | TGTTGGTTTACCGTTCGGTG |
|  | *pduU-R* | | TGGGGCCGATAAACATCAAAC |
| ***pduV*** | *pduV-F* | | TTTCTCGATCGCTTTACCGG |
|  | *pduV-R* | | TAATTGTCACCTGCGCATCT |

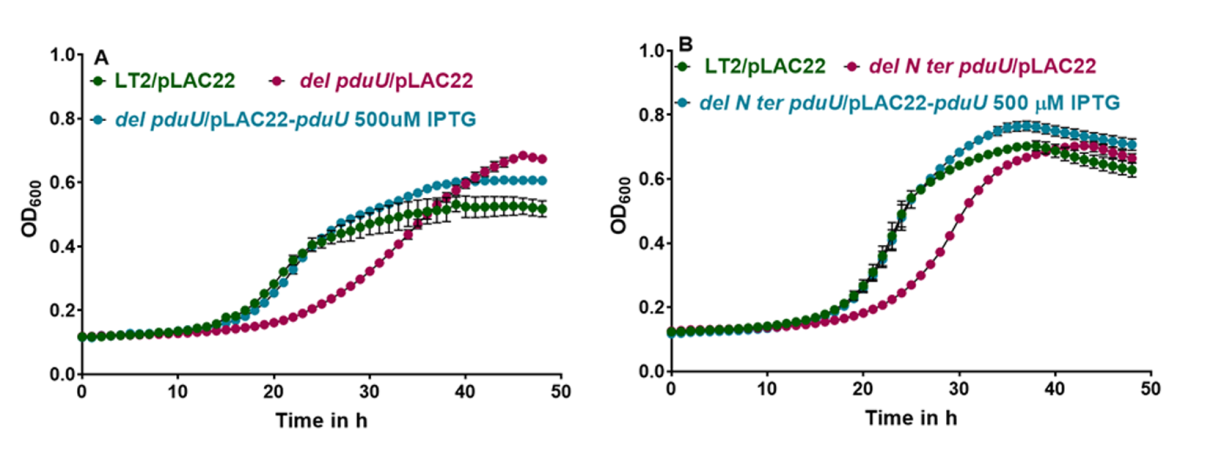

**Fig S1:** Complementation of *del pduU* (A) and *del* *N-ter pduU* (B) by ectopic expression through pLAC22-*pduU* at 500 µM IPTG. The error bars represent one standard deviation and are based on three biological replicates.

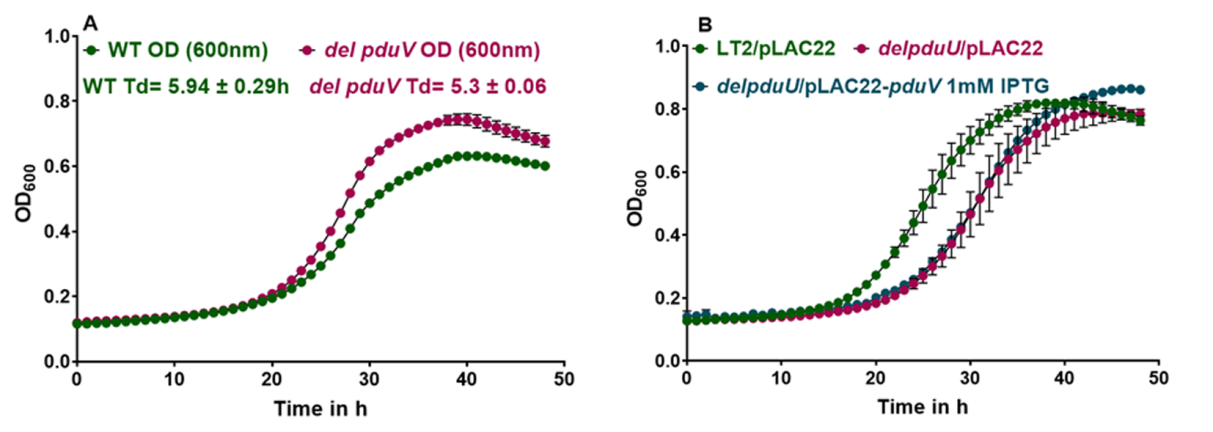

**Fig S2:** A: The growth of *del pduV* mutant at 37 ºC in minimal medium supplemented with 0.4% 1,2 propanediol as carbon source with saturating B_12_ (100nM) concentration. B: Cross-complementation of a *del pduU* mutant by ectopic expression of pLAC22- *pduV.* The error bars represent one standard deviation and are based on three biological replicates.

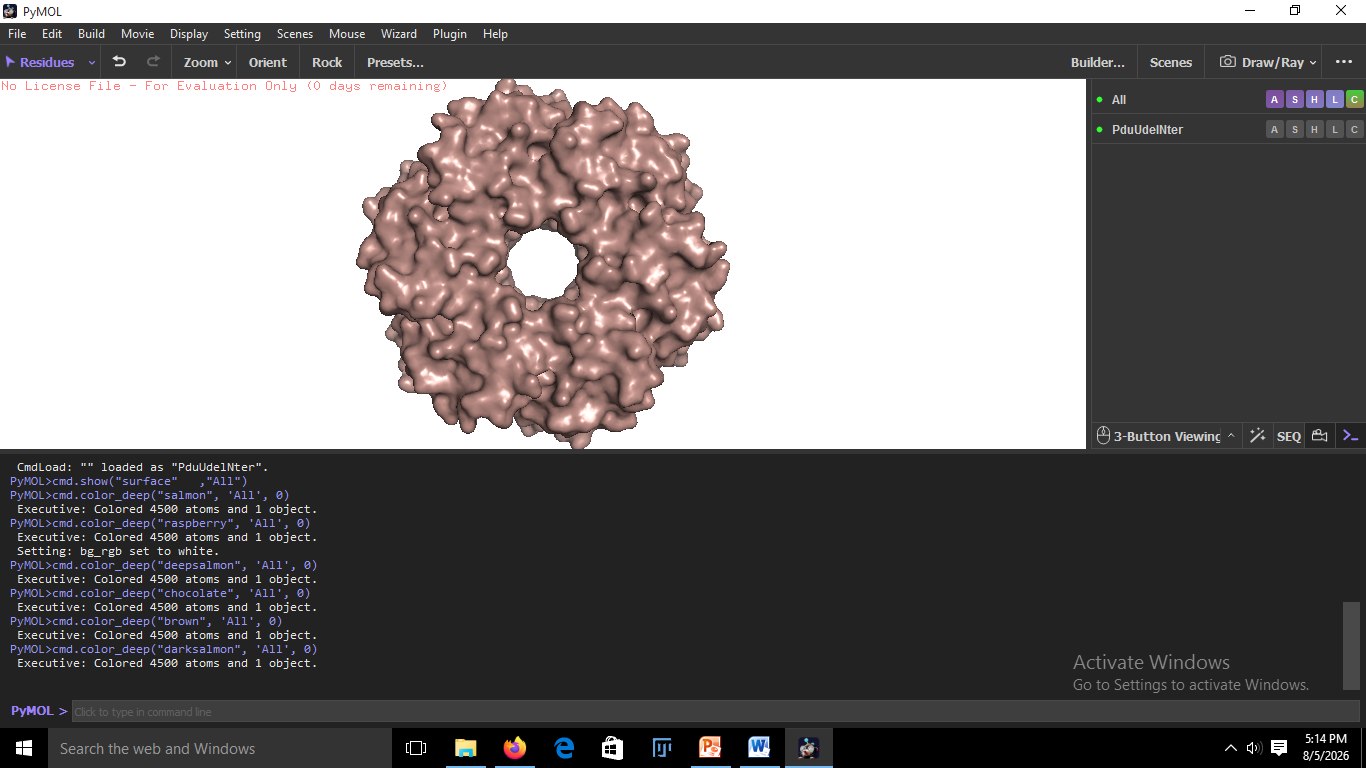

**13.5 Å**

**PduU N-ter deletion model**

**Propionic acid**

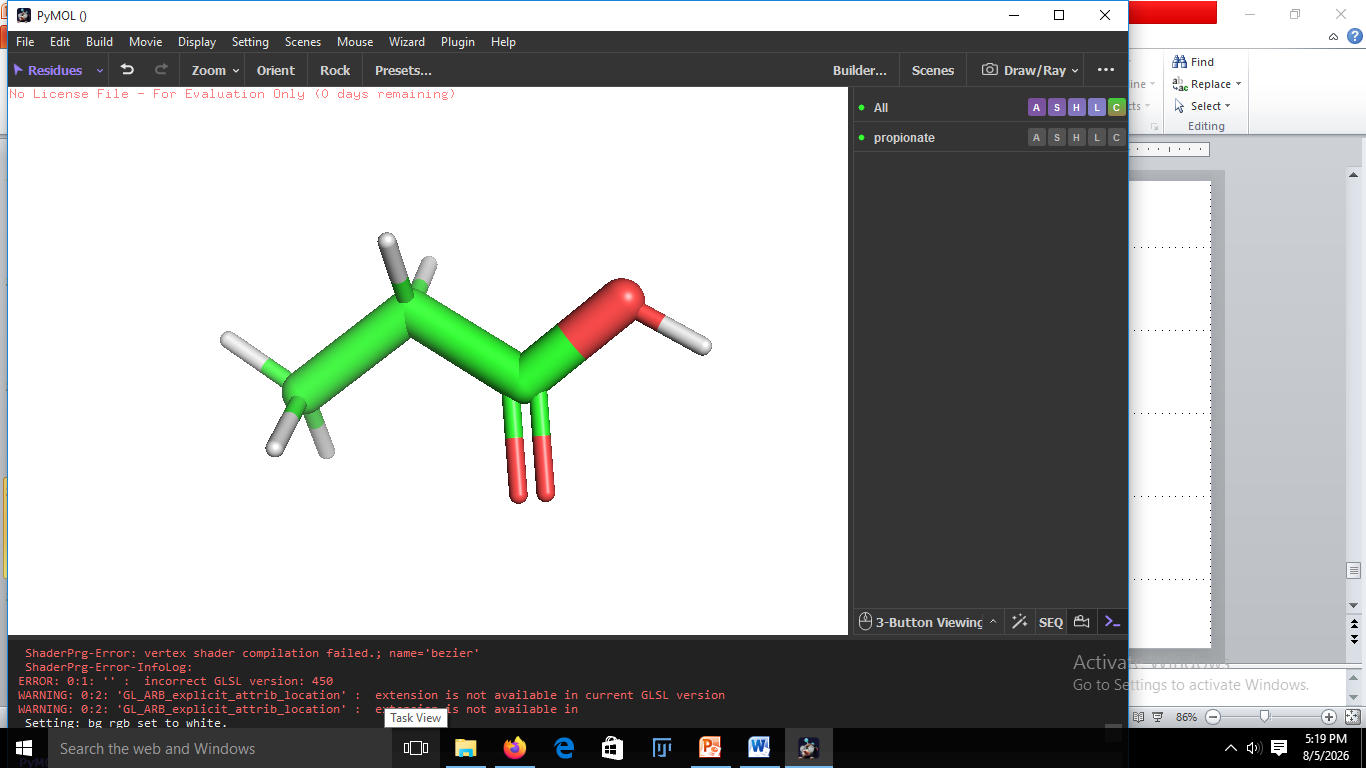

**5.3 Å**

**1.8 Å**

**Fig S3:** Structural modelling of the PduU shell protein using the Swiss-Model platform, with the N-terminal β-barrel domain truncated revealed an open central pore measuring 13.5. PyMOL-based molecular dimension analysis showed that propionate exhibits a maximum longitudinal length of approximately 5.8 Å and a cross-sectional width of 1.8 Å Given that the molecular dimensions of propionate are substantially smaller than the 13.5 Å internal pore aperture, the truncated PduU variant establishes a sterically unobstructed pore capable of facilitating the unhindered flux of propionate across the microcompartment shell.
